# Cryo-EM Structure of the Photosynthetic LH1-RC Complex from *Rhodospirillum rubrum*

**DOI:** 10.1101/2021.05.30.446358

**Authors:** K. Tani, R. Kanno, X.-C. Ji, M. Hall, L.-J. Yu, Y. Kimura, M. T. Madigan, A. Mizoguchi, B. M. Humbel, Z.-Y. Wang-Otomo

**Author notes:** These authors contributed equally to this work: K. Tani, R. Kanno.

## Abstract

We present a cryo-EM structure of the light-harvesting-reaction center (LH1-RC) core complex from purple phototrophic bacterium *Rhodospirillum* (*Rsp*.) *rubrum* at 2.76 Å resolution. The LH1 complex forms a closed, slightly elliptical ring structure with 16 αβ-polypeptides surrounding the RC. Our biochemical analysis detected rhodoquinone (RQ) molecules in the purified LH1-RC, and the cryo-EM density map specifically positions RQ at the Q_A_ site in the RC. The geranylgeraniol sidechains of bacteriochlorophyll (BChl) *a*_G_ coordinated by LH1 β-polypeptides exhibit a highly homologous tail-up conformation that allows for interactions with the bacteriochlorin rings of nearby LH1 α-associated BChls *a*_G_. The structure also revealed key protein–protein interactions in both N- and C-terminal regions of the LH1 αβ-polypeptides, mainly within a face-to-face structural subunit. Our findings enable to evaluate past experimental and computational results obtained with this widely used organism and provide crucial information for more detailed exploration of light-energy conversion, quinone transport, and structure–function relationships in pigment-protein complexes.

---

*Rsp. rubrum* is an anoxygenic phototrophic purple bacterium with a long history as a model for the study of bacterial photosynthesis and related metabolic processes. It is unique among purple bacteria by producing both rhodoquinone (RQ) and ubiquinone (UQ)^*1*^ as electron carriers and bacteriochlorophyll (BChl) *a* esterified at the propionic acid sidechain by geranylgeraniol (abbreviated as BChl *a*_G_) rather than phytol^*2*^. *Rsp. rubrum* has a single pair of αβ-polypeptides in its core light-harvesting (LH1) complex and lacks both the peripheral light-harvesting (LH2) complex and reaction center (RC) cytochrome (Cyt) *c* subunit present in many purple bacteria; thus, *Rsp. rubrum* is one of the simplest phototrophic bacteria known, in terms of its photosynthetic light reactions. Despite many efforts, structures of both purified LH1 and the RC-associated core complex (LH1-RC) of *Rsp. rubrum* have not been obtained at high resolution, and no RC atomic structure is known.

Here we present a cryo-EM structure of the *Rsp. rubrum* LH1-RC complex at 2.76 Å resolution (*Supplementary* Table S1, Figs. S1–3). The LH1 complex forms a closed, slightly elliptical double ring composed of 16 pairs of α(inner)β(outer)-polypeptides, 32 BChls *a*_G_ and 16 *all*-*trans*-spirilloxanthins (Fig. 1). The arrangement of *Rsp. rubrum* LH1 α- and β-polypeptides overlaps with that of the LH1 from the thermophilic purple sulfur bacterium *Thermochromatium* (*Tch*.) *tepidum* (Fig. 1d)^*3*^. The N-terminal Met in the LH1 α-polypeptide is formylated (*Supplementary* Fig. S4)^*4*^, and the formyl groups were modeled in our structure. Both the LH1 α- and β-polypeptides of *Rsp. rubrum* exhibit helical conformations in the transmembrane regions, consistent with those observed by solution NMR (*Supplementary* Fig. S5)^*5*^. The *Rsp. rubrum* RC is surrounded by the LH1 complex with only a few close contacts on the periplasmic surface with residues near Ser34 in the LH1 α-polypeptides. There are no apparent strong interactions of the *Rsp. rubrum* RC with its LH1 polypeptides as occurs in Cyt *c*-bound LH1-RCs where the C-terminal domains of some LH1 α-polypeptides interact extensively with the Cyt *c* subunit^*6*^. This may explain why an LH1-only complex can be easily purified from *Rsp. rubrum* (*Supplementary* Fig. S6a)^*7*^. Cofactors in the *Rsp. rubrum* RC include four BChls *a*_G_, two bacteriopheophytins (BPhe) *a*_G_, one 15-*cis*-spirilloxanthin, an RQ-10 at the Q_A_ site, and a UQ-10 at the Q_B_ site (Fig. 1c).

**Fig. 1.**
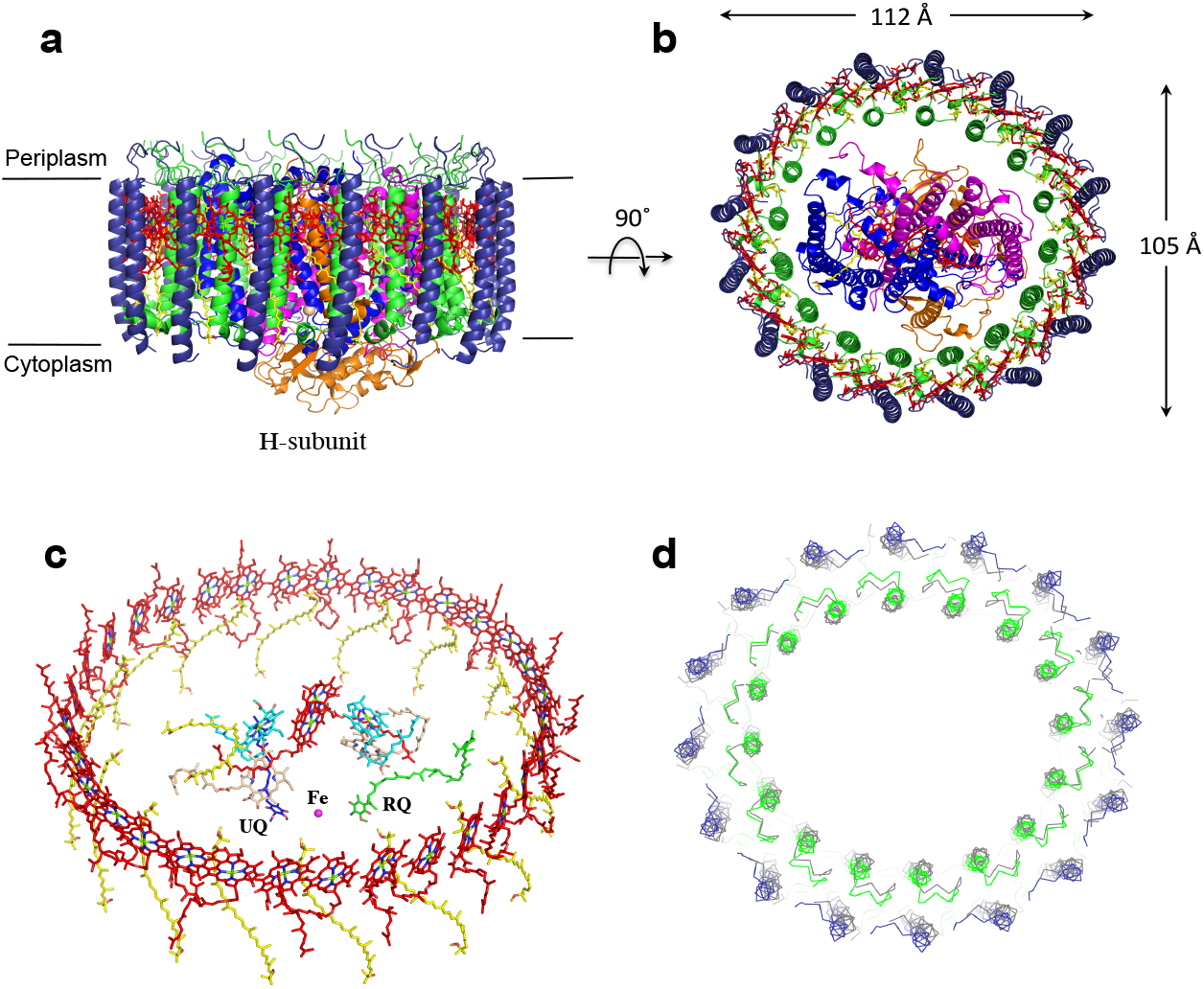
Structure overview of the *Rsp. rubrum* LH1-RC complex. **(a)** Side view of the LH1-RC parallel to the membrane plane. **(b)** Top view of the LH1-RC from the periplasmic side of the membrane. **(c)** Tilted view of the cofactor arrangement. **(d)** Superposition of Cα carbons of the LH1 αβ-polypeptides between *Rsp. rubrum* and *Tch. tepidum* (gray, PDB: 5Y5S). Color scheme: LH1-α, green; LH1-β, slate-blue; L-subunit, magenta; M-subunit, blue; BChl *a*_G_ in LH1 and special pair, red sticks; Accessory BChl *a*_G_, cyan sticks; BPhe *a*_G_, light-pink sticks; Spirilloxanthin, yellow sticks; UQ-10, blue sticks; RQ-10, green sticks; Fe, magenta ball. Phospholipids and detergents are omitted for clarity.

The BChl *a*_G_ molecules in *Rsp. rubrum* LH1 form an elliptical, partially overlapping ring with average Mg–Mg distances of 9.3 Å within a dimer and 8.5 Å between dimers (Fig. 1c). These molecules are ligated by histidine residues (α-His29 and β-His39, Fig. 2) that have His–Mg distances of 2.3 Å and 2.0 Å, respectively (*Supplementary* Table S2). The geranylgeranyl sidechains in the BChl *a*_G_ associated with β-polypeptides form a tail-up conformation (Fig. 2a) with a much higher structural homogeneity compared with those of purple bacteria whose BChl *a* is esterified by a phytyl group (*Supplementary* Fig. S7). This may be due to the multiple double bonds present in the geranylgeranyl group, which results in a more rigid conformation. This unique conformation allows the geranylgeranyl sidechains of the β-associated BChl *a*_G_ to interact with the bacteriochlorin ring of the α-associated BChl *a*_G_ (Fig. 2b) where the methyl group at geranylgeranyl g11 points to the macrocycle of a neighboring BChl *a*_G_ at a distance of ∼3.5 Å. This close proximity likely allows for π-π interactions between the double bonds in the geranylgeranyl group and nearby bacteriochlorins. The C3-acetyl oxygen atoms in BChl *a*_G_ form hydrogen bonds with the Trp residues (α-Trp40 and β-Trp48) of their associated polypeptides (Figs. 2c and 2d), in agreement with the results of resonance Raman spectroscopy^*8-10*^. The aromatic side chains of the Trp residues also interact with bacteriochlorins through π-π stacking, further stabilizing the LH1 complex.

**Fig. 2.**
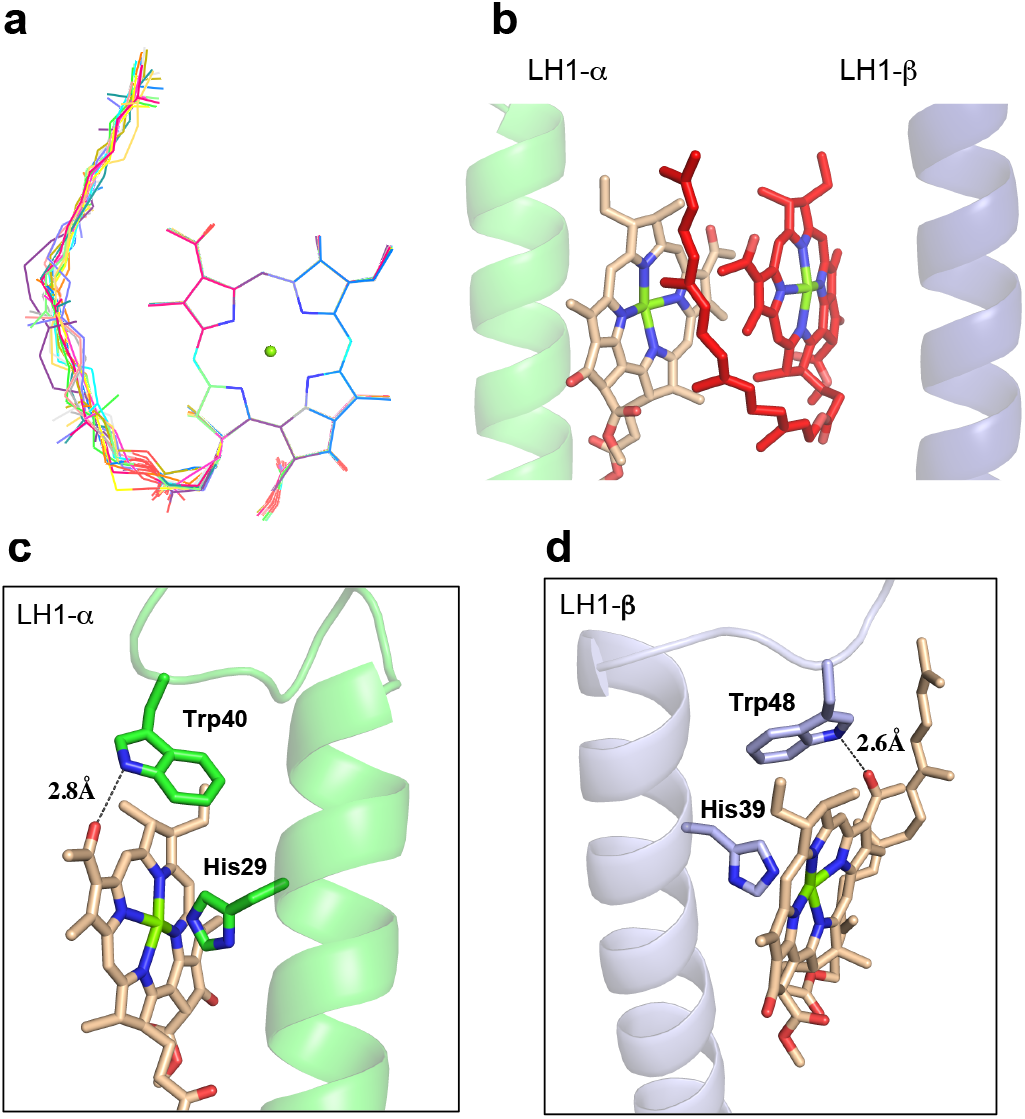
BChl *a*_G_ in *Rsp. rubrum* LH1 complex. **(a)** Superposition of the bacteriochlorin rings of 16 BChl *a*_G_ molecules bound to the LH1 β-polypeptides. **(b)** Interactions between a geranylgeranyl sidechain of the BChl *a*_G_ (red sticks) bound to LH1 β-polypeptide and the bacteriochlorin ring (tint sticks) of a BChl *a*_G_ bound to the LH1 α-polypeptide. **(c)** Coordination and hydrogen bonding of the BChl *a*_G_ in LH1 α-polypeptide. **(d)** Coordination and hydrogen bonding of the BChl *a*_G_ in LH1 β-polypeptide.

The *Rsp. rubrum* RC is composed of L, M and H subunits with an overall protein structure and cofactor organization as reported in other purple bacteria (Figs. 3a and 3b). Our biochemical analyses of purified LH1-RC complexes using HPLC, absorption, and MALDI-TOF/MS spectroscopies identified approximately one UQ-9, ten UQ-10 and one RQ-10 to be present per complex (Fig. 3c, *Supplementary* Fig. S8). The cryo-EM density map at the Q_A_ site is best modeled by the head group of an RQ-10 (Fig. 3d, *Supplementary* Fig. S3b). The aminobenzoquinone ring of RQ-10 is stabilized by formation of two hydrogen bonds between its carbonyl groups and a His residue (M-His218) and mainchain amide group (M-Ala259) in a loop region of the M-subunit (*Supplementary* Fig. S9). By contrast, the density map of UQ-10 at the Q_B_ site was more diffuse (*Supplementary* Fig. S3b), indicating a more mobile nature of this quinone. The special pair BChl *a*_G_ molecules in the *Rsp. rubrum* RC are ligated by histidine residues (L-His174 and M-His201) with His–Mg distances (2.09 Å and 2.12 Å, respectively) and BChl *a*_G_–BChl *a*_G_ distance (7.76 Å) similar to those of *Tch. tepidum* (*Supplementary* Table S2, Fig. S10). While the C3-acetyl group in the *Rsp. rubrum* special pair L-BChl *a*_G_ forms a hydrogen bond with a His residue (L-His169) as in all other purple bacterial RCs, no such hydrogen bond is possible to the C3-acetyl group of M-BChl *a*_G_ because the nearby residue is a Phe (M-Phe196) (*Supplementary* Fig. S10), as is also true of the RC from *Rhodobacter* (*Rba*.) *sphaeroides*. The corresponding residue is Tyr in RCs from the purple bacteria *Blastochloris viridis, Tch. tepidum, Thiorhodovibrio* strain 970 and *Rhodopseudomonas palustris* that also form a hydrogen bond with the M-BChl *a* C3-acetyl group.

**Fig. 3.**
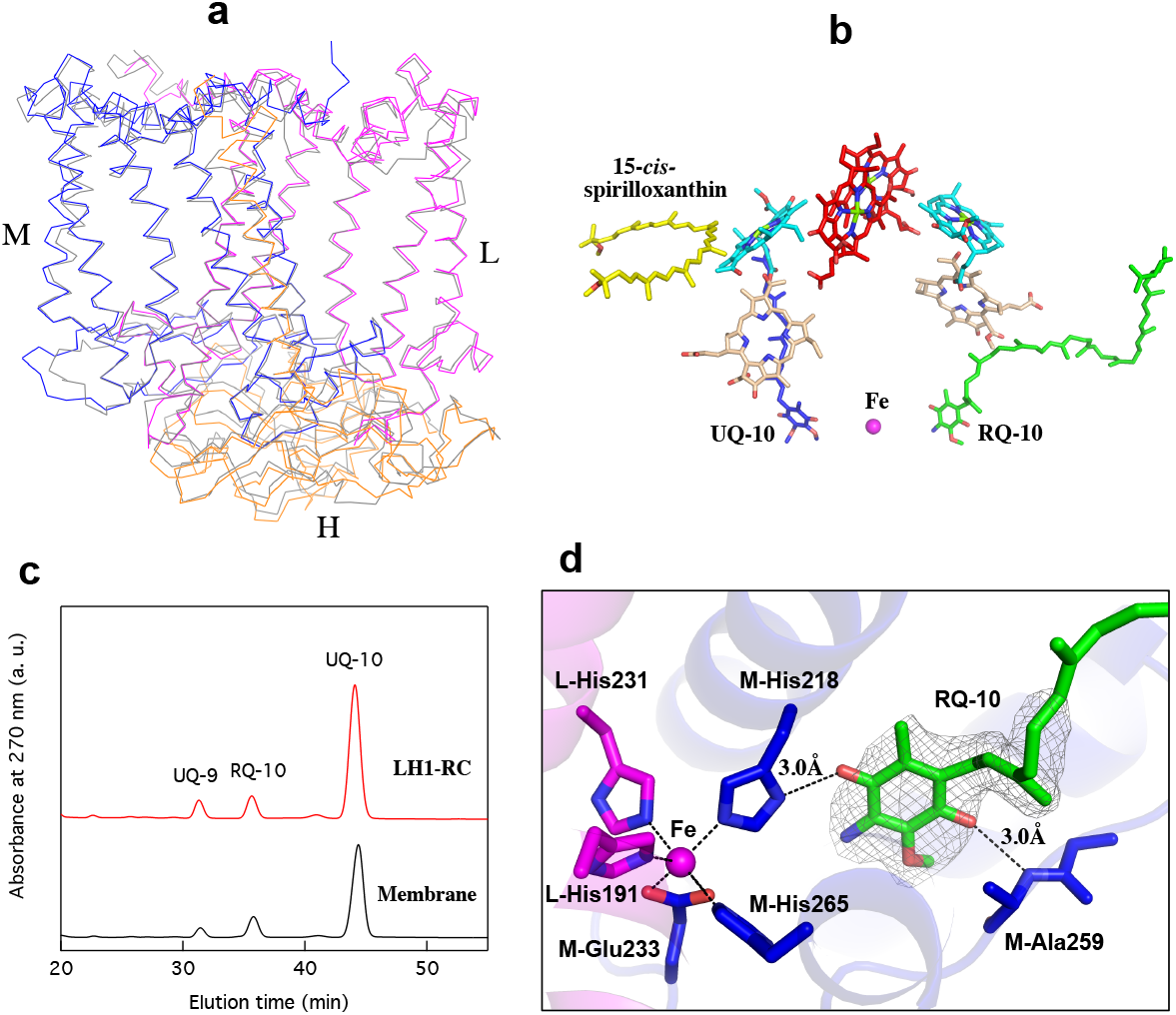
The RC complex of *Rsp. rubrum*. **(a)** Side view of superposition of Cα carbons for the RC protein subunits between *Rsp. rubrum* (colored) and *Rba. sphaeroides* (gray, PDB: 3I4D). **(b)** Cofactor arrangement. Color code: special pair BChl *a*_G_, red; accessory BChl *a*_G_, cyan; BPhe *a*_G_, light-pink; 15-*cis*-spirilloxanthin, yellow; UQ-10, blue; RQ-10, green; Fe, magenta. **(c)** Reverse-phase HPLC chromatograms of the quinones extracted from purified LH1-RC and membranes. **(d)** The RQ-10 modeled at Q_A_ site and its interactions with the surrounding residues in the RC. Density map for the RQ-10 head group is shown at contour level of 4σ.

A total of 10 phospholipids (2 PG, 5 CL and 3 PE) were modeled in the cavities between the RC and LH1 based on the density map (Fig. 4a and 4b). Most CL molecules were located on the cytoplasmic side of the membrane with their head groups pointing toward the membrane surface, whereas most PG and PE molecules were distributed on the periplasmic side. This phospholipid distribution is similar to that observed in other LH1-RCs^*3, 6, 11, 12*^. Channels were found between every adjacent pair of LH1 αβ-polypeptides (Fig. 4c and 4d) and are composed of nine hydrophobic residues in each polypeptide (*Supplementary* Fig. S4). Similar channels have also been observed in other LH1 complexes with closed ring-like structures^*3, 6, 12*^, suggesting that these hydrophobic channels function as paths for quinone transport into and out of the complex. Such a function is supported by molecular dynamics simulation^*13*^ and biochemical analysis^*14*^.

**Fig. 4.**
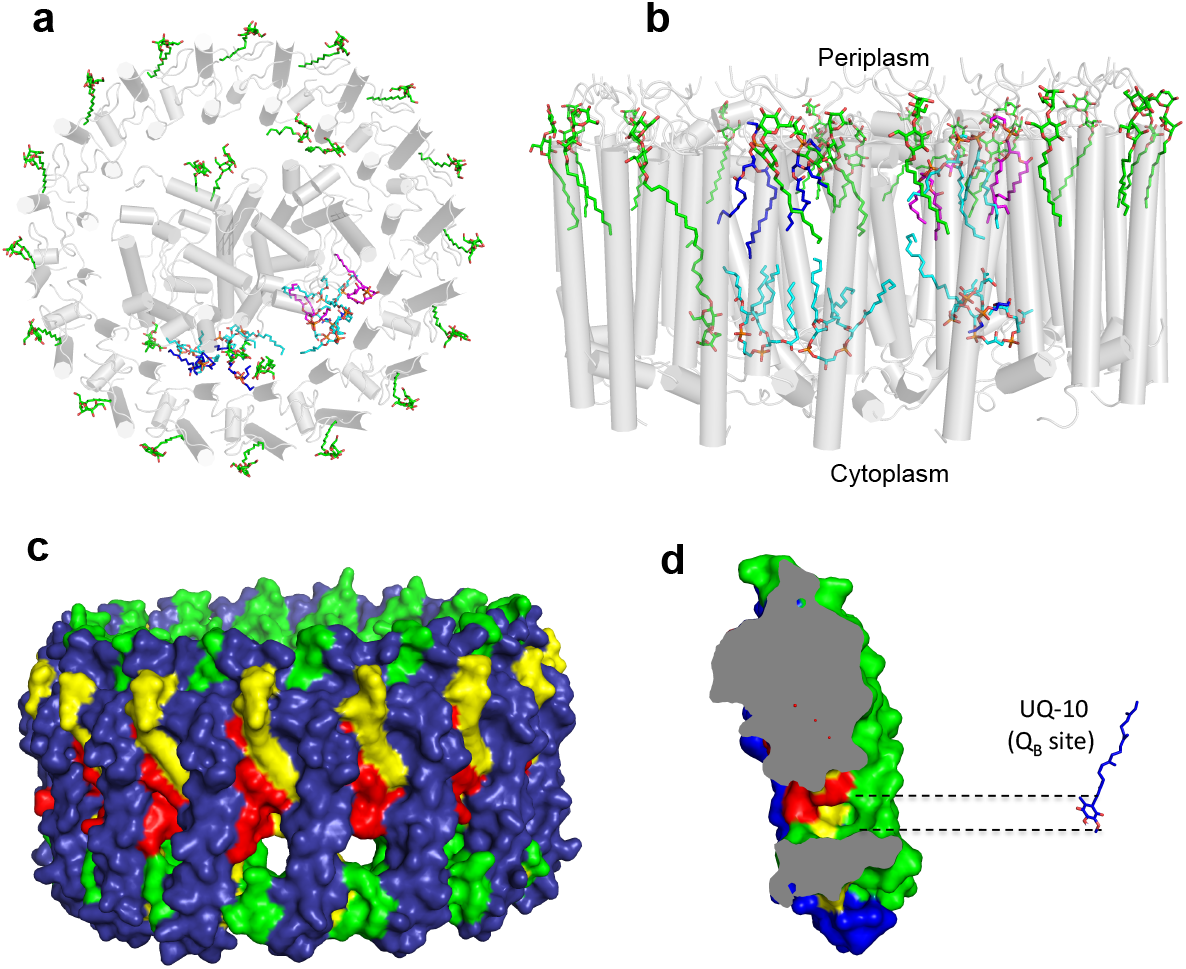
Phospholipids, detergents and channels in LH1-RC complex. Top view **(a)** and side view **(b)** of the phospholipid and detergent distributions for CL (cyan), PG (magenta), PE (blue) and DDM (green). All proteins are shown in gray. **(c)** Surface representation of the channels in LH1 ring with color codes: LH1-α, green; LH1-β, slate-blue; BChl *a*_G_, red; spirilloxanthin, yellow. **(d)** Cross section of a putative quinone channel illustrating the channel opening and the size of the benzoquinone head group of the UQ-10 (blue sticks) at Q_B_ site.

The *Rsp. rubrum* LH1 complex is particularly well known for its ability to form a highly stable structural subunit with an absorption maximum at 820 nm (*Supplementary* Fig. S6a)^*15*^. This so-called B820 subunit can be obtained from the native LH1 complex at high detergent concentrations or reconstituted from individually separated polypeptides including those from other species^*16-18*^ and various BChl analogues^*19, 20*^. The B820 subunit was determined by spectroscopic^*21, 22*^, biochemical^*23, 24*^ and small-angle neutron scattering^*25*^ methods to be a heterodimer composed of two interacting BChl *a* molecules linked to one αβ-polypeptide pair. Two different subunit forms can be distinguished from the LH1 complex as shown in Fig. 5a, one with a face-to-face configuration and another with a back-to-back configuration for the bacteriochlorins. Solution NMR^*26*^ and reconstitution^*27*^ experiments established that the B820 subunit has the face-to-face configuration with π-overlap at pyrrole rings III and V.

**Fig. 5.**
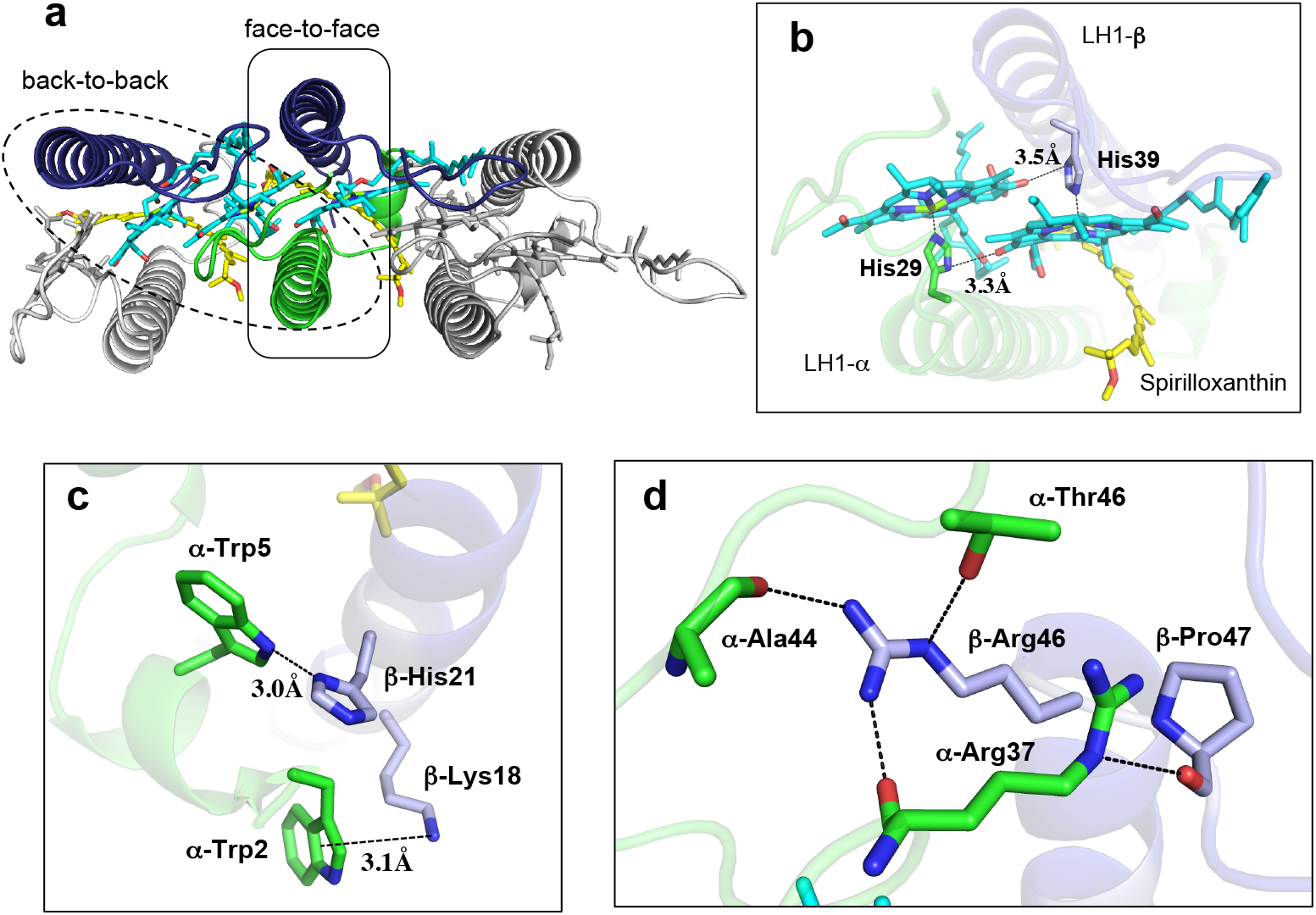
Structural subunits in the LH1 complex. **(a)** Two structural subunits with different configurations for the BChl *a*_G_ dimers. Color codes in the subunits: LH1-α, green; LH1-β, slate-blue; BChl *a*_G_, cyan; spirilloxanthin, yellow. Other polypeptides and pigments are shown in gray. **(b)** Structure of the face-to-face subunit showing coordination and cross hydrogen-bonding of the His residues to BChl *a*_G_ molecules. **(c)** Major interactions in the N-terminal region within a face-to-face subunit. **(d)** Major interactions in the C-terminal region within a face-to-face subunit. Dashed lines indicate distances shorter than 3.5 Å.

Due to its structural simplicity and flexibility, the *Rsp. rubrum* B820 subunit has been thoroughly investigated using protease-modified polypeptides, mutants and chemically synthesized polypeptides^*27-30*^, the results from which predicted minimal requirements for stabilizing the subunit structure. These requirements included (i) a central α-helical transmembrane domain composed of 18 hydrophobic residues; (ii) a His residue for coordination and hydrogen bonding to different BChl molecules; (iii) a Trp residue for hydrogen bonding to the BChl C3^1^ carbonyl oxygen; and (iv) N-terminal regions of α- and β-polypeptides. The His interaction was estimated to account for over half of the stabilization energy of the B820 subunit, followed by hydrogen bonding by Trp residues^*30*^. Now, these predictions can be modeled by our high-resolution *Rsp. rubrum* LH1 structure. Both α- and β-polypeptides form transmembrane α-helices in the regions of 12–34 and 19–42, respectively (Fig. 1a, *Supplementary* Fig. S4). The BChl *a*-coordinating His residue also forms a hydrogen bond with the C13^1^ carbonyl oxygen atom of the neighboring BChl *a*, and this cross-hydrogen bonding only occurs within the face-to-face subunit (Fig. 5b). Both Trp residues (α-Trp40 and β-Trp48) form hydrogen bonds with the C3-acetyl oxygen atoms of BChl *a* as was shown in Figs. 2c and 2d.

In addition to these pigment–protein interactions, our structure revealed significant protein– protein interactions in the N- and C-terminal domains of the *Rsp. rubrum* LH1 subunit. All amino acids at the N-terminal end of the α-polypeptide including their formyl groups can be traced in our cryo-EM density map, indicating a stable, ordered conformation (*Supplementary* Fig. S3b). The α-Trp5 forms a relatively strong hydrogen bond with β-His21, and α-Trp2 likely employs cation–π interactions^*31*^ with β-Lys18 in the N-terminal region (Fig. 5c). The α-Trp5 and β-His21 residues are conserved in all LH1 complexes reported, and the hydrogen bond between them has been observed in the LH1 structures of *Tch. tepidum*^*3*^, *Trv*. strain 970^*6*^ and *Rps. palustris*^*12*^. The α-Trp2 and β-Lys18 are either conserved or replaced with an aromatic (Tyr, His) and a cationic (Arg) residue, respectively, at corresponding positions in many LH1 complexes (*Supplementary* Fig. S4). Our *Rsp. rubrum* LH1 structure also explains previous observations that a stretch of 9–13 amino acids at the N-terminal end of the α- and β-polypeptides (especially the first Met of the α-polypeptide) is important for stabilizing the B820 subunit^*4, 27*^ and that the first three residues of the α-polypeptide likely maintain close interactions with the sidechain of His21 of the β-polypeptide^*27*^.

In summary, our high-resolution cryo-EM structure of the *Rsp. rubrum* LH1-RC definitively locates components of the complex with respect to each another, and reveals their intimate interactions and dependencies, thus providing a blueprint from which an even deeper understanding of light-energy conversion and quinone transport in photosynthetic organisms can be pursued. Our work also provides clues for how structure confers stability in pigment-protein complexes and broadens our understanding of the spectroscopy, photochemistry, and computational biochemistry of both the entire *Rsp. rubrum* LH1-RC complex and its simplified B820 subunit.

## ACKNOWLEDGMENT

This research was partially supported by Platform Project for Supporting Drug Discovery and Life Science Research (Basis for Supporting Innovative Drug Discovery and Life Science Research (BINDS)) from AMED under Grant Numbers JP21am0101118 (support number 1758) and JP21am0101116 (support number 1878), 17am0101116, 18am0101116, 19am0101116 and 20am0101116. R.K., M.H. and B.M.H. acknowledge the generous support of the Okinawa Institute of Science and Technology and the Japanese Cabinet Office. This work was supported in part by JSPS KAKENHI Grant Numbers JP16H04174, JP18H05153, JP20H05086 and JP20H02856, Takeda Science Foundation, and the Kurata Memorial Hitachi Science and Technology Foundation, Japan, and the National Key R&D Program of China (No. 2019YFA0904600).

## Author Contributions

Z.-Y.W.-O. and K.T. designed the work, K.T., R.K., X.-C. J. and M.H. performed the experiments, K.T., R.K., L.-J.Y., Y.K., M.T.M., A.M., B.M.H. and Z.-Y.W.-O. analyzed data, Z.-Y.W.-O., K. T. and M.T.M. wrote the manuscript. All authors edited and revised the manuscript.

## Data availability

Map and model have been deposited in the EMDB and PDB with the accession codes: EMD-31258 and PDB-7EQD. All other data are available from the authors upon reasonable request.

