## Supplementary Information for "Cryo-EM Structure of the Photosynthetic LH1-RC Complex from *Rhodospirillum rubrum*"

### Materials and Methods

**Preparation and Characterization of the LH1-RC Complex.** *Rsp. rubrum* NBRC3986 (type strain, Esmarch 1887, Molisch 1907) was purchased from National Institute of Technology and Evaluation, Japan. The cells were cultivated phototrophically (anoxic/light) at room temperature for 7 days under incandescent light (60W). Preparation of the *Rsp. rubrum* LH1-RC was conducted by solubilizing chromatophores with 1.0 % w/v DDM in 20 mM Tris-HCl (pH 8.5) buffer for 60 min at room temperature, followed by differential centrifugation. The supernatant was loaded onto a DEAE column (Toyopearl 650S, TOSOH) equilibrated at 4 ˚C with 20 mM Tris-HCl buffer (pH 7.5) containing 0.1 % w/v of DDM. The LH1-RC fraction was eluted by a linear gradient of $CaCl_2$ from 0 mM to 100 mM. The peak fractions with $A_{877}/A_{280} >$ 2.2 were collected for subsequent measurement (*Supplementary* Fig. S6)[1], and then assessed by negative-stain EM using a JEM-1011 instrument (JEOL) (*Supplementary* Fig. S1a). Quinones were extracted from the purified LH1-RC and chromatophores, and quinone contents were analyzed (*Supplementary* Fig. S8) using the methods described previously[2]. Circular dichroism (CD) and magnetic CD spectra were recorded on a Jasco J-720w spectropolarimeter in the range from 400 nm to 1000 nm under same conditions as those given before[3].

**Cryo-EM Data Collection.** Proteins for cryo-EM were concentrated to 3.5 mg/ml. Three microliters of the protein solution were applied on a glow-discharged holey carbon grids (200 mesh Quantifoil R2/2 molybdenum), which had been treated with $H_2$ and $O_2$ mixtures in a Solarus plasma cleaner (Gatan, Pleasanton, USA) for 30 s and then blotted, and plunged into liquid ethane at –182 ˚C using an EM GP2 plunger (Leica, Microsystems, Vienna, Austria). The applied parameters were a blotting time of 5 s at 80% humidity and 4°C. Data were collected at OIST on a Titan Krios (Thermo Fisher Scientific, Hillsboro, USA) electron microscope at 300 kV equipped with a Falcon 3 camera (Thermo Fisher Scientific). Movies were recorded using EPU software (Thermo Fisher Scientific) at a nominal magnification of 96 k in counting mode and a pixel size of 0.820 Å with a dose rate of 0.92 e- per physical pixel per second, corresponding to 1.37 e- per $Å^2$ per second at the specimen level. The exposure time was 30.6 s, resulting in an accumulated dose of 42 e- per $Å^2$. Each movie includes 40 fractioned frames. Pixel size calibration was done with a known protein structure determined by X-ray crystallography.

**Image Processing.** All of the stacked frames were subjected to motion correction with MotionCor2[4]. Defocus was estimated using CTFFIND4[5]. A total of 262,517 particles were selected from 1,701 micrographs using the EMAN2 suite (*Supplementary* Fig. S2a)[6]. The particles were further analyzed with RELION3.1[7], and 155,557 particles were selected by 2-D classification. At first, the initial 3-D models were generated from these particles using EMAN2 or RELION3.1[7], but the 3D-reconstruction starting from either model could not reach beyond 4 Å resolution despite obtaining good 2D class average (*Supplementary* Fig. S1c). We then tried to build the initial model from the cryo-EM map of *Trv.* strain 970 LH1-RC[8] after subtraction of the density corresponding to the C-subunit and the C-terminal portions of LH1 α- and β-subunits. Supervised 3-D classification of the 155,557 particles were performed using the modified *Trv.* strain 970 LH1-RC cryo-EM map as initial model resulting in two good classes containing 145,033 particles. The 3-D auto refinement without any imposed symmetry (C1) produced a map at 2.86 Å resolution after contrast transfer function refinement, Bayesian polishing, masking, and post-processing. All of the particle projections were subjected to subtraction of the detergent micelle density followed by 3-D auto refinement to yield the final map with a resolution of 2.76 Å according to the gold-standard Fourier shell correlation using a criterion of 0.143 (*Supplementary* Fig. S2b)[9]. The local resolution maps were calculated on RESMAP[10].

**Model Building and Refinement of the LH1-RC Complex.** The atomic model of the *Tch. tepidum* LH1-RC (PDB: 5Y5S) was fitted to the cryo-EM map obtained for the *Rsp. rubrum* LH1-RC using Chimera[11]. Amino acid substitutions and real space refinement for the peptides and cofactors were performed using COOT[12]. The C-terminal regions of the LH1 αβ-subunit were modelled *ab-initio* based on their density. The manually modified model was real space refined on PHENIX[13], and the COOT/PHENIX refinement was iterated until the refinements converged. The statistics were calculated using MolProbity[14]. Figures were drawn with the Pymol Molecular Graphic System (Schrödinger)[15] and UCSF Chimera[11].

**Table S1 Cryo-EM data collection, refinement and validation statistics of the *Rsp. rubrum* LH1-RC complex**

| | LH1-RC complex<br>(EMDB-31258)<br>(PDB-7EQD) |
|---|---|
| **Data collection and processing** | |
| Magnification | 96000 |
| Voltage (kV) | 300 |
| Electron exposure (e–/Å$^2$) | 38 |
| Defocus range (μm) | -0.9 to -3.2 |
| Pixel size (Å) | 0.820 |
| Symmetry imposed | C1 |
| Initial particle images (no.) | 262517 |
| Final particle images (no.) | 1450334 |
| Map resolution (Å) | 2.8 |
| FSC threshold | 0.143 |
| Map resolution range (Å) | 295-2.8 |
| **Refinement** | |
| Initial model used (PDB code) | 5Y5S |
| Model resolution (Å) | 2.9 |
| FSC threshold | 0.5 |
| Model resolution range (Å) | 133-2.8 |
| Map sharpening *B* factor (Å$^2$) | -65 |
| Model composition | |
| Non-hydrogen atoms | 23122 |
| Protein residues | 2302 |
| Ligands | 92 |
| *B* factors (Å$^2$) | |
| Protein | 36.2 |
| Ligand | 26.4 |
| R.m.s. deviations | |
| Bond lengths (Å) | 0.005 |
| Bond angles (°) | 1.013 |
| Validation | |
| MolProbity score | 1.75 |
| Clashscore | 10.60 |
| Poor rotamers (%) | 0.05 |
| Ramachandran plot | |
| Favored (%) | 96.71 |
| Allowed (%) | 3.29 |
| Disallowed (%) | 0.00 |

**Table S2 Comparison of the distances of His–BChl(Mg) and BChl(Mg)–BChl(Mg) in LH1, LH2 and RC special pairs from various phototrophic bacteria.**

| | Distance of His(Nε2) to BChl–Mg (Å)[a] | | Distance of Mg–Mg (Å)[a] | |
|---|---|---|---|---|
| LH1 or LH2 | α | β | Long | Short |
| ***Rsp. rubrum* (LH1)** | **2.27** | **2.03** | **9.34** | **8.51** |
| *Rps. palustris* (LH1-W) | 2.93 | 2.71 | 9.61 | 8.29 |
| *Tch. tepidum* (LH1) | 2.19 | 2.19 | 8.88 | 8.72 |
| *Trv.* strain 970 (LH1) | 2.33 | 2.31 | 8.90 | 8.46 |
| *Blc. viridis* (LH1) | 2.54 | 2.25 | 8.8 | 8.5 |
| *Rfx. castenholzii* (B880) | 2.32 | 2.29 | 9.5 | 9.3 |
| *Rps. acidophila* (B850) | 2.34 | 2.34 | 9.5 | 8.8 |
| *Phs. molischianum* (B850) | 2.27 | 2.32 | 9.2 | 8.9 |
| RC (special pair) | L-subunit | M-subunit | BChl *a* (L)–BChl *a* (M) | |
| ***Rsp. rubrum*** | **2.09** | **2.12** | **7.76** | |
| *Rba. sphaeroides* | 2.27 | 2.06 | 7.84 | |
| *Rps. palustris* | 2.73 | 2.74 | 7.69 | |
| *Tch. tepidum* | 2.17 | 2.19 | 7.87 | |
| *Trv.* strain 970 | 2.33 | 2.31 | 7.65 | |
| *Blc. viridis* | 2.36 | 2.35 | 7.83 | |

[a] These values were derived from Protein Data Bank: 5Y5S for *Tch. tepidum* LH1-RC, 6ET5 for *Blc. viridis* LH1-RC, 5YQ7 for *Rfx. castenholzii* B880, 1NKZ for *Rps. acidophila* LH2, 1LGH for *Phs. molischianum* LH2, 2J8C for *Rba. sphaeroides* RC.

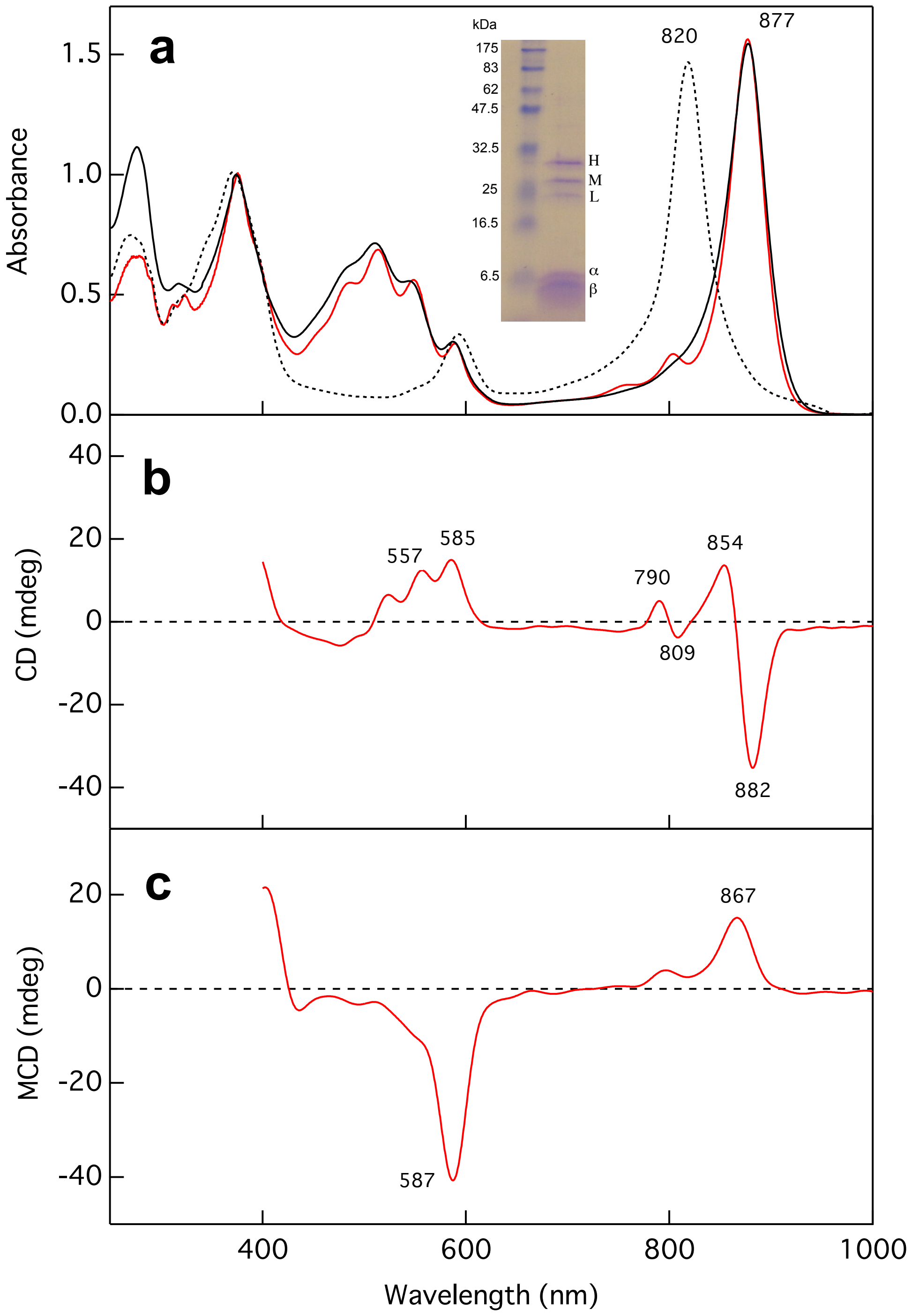


**Fig. S1 Absorption (a), circular dichroism (b) and magnetic circular dichroism (c) spectra of the purified *Rsp. rubrum* LH1-RC complex (red solid curves) at room temperature.** Absorption spectra of the LH1-only complex and a B820 subunit are shown by black solid and dotted curves, respectively. Inset: Coomassie blue-stained 12% SDS-PAGE gel for the purified LH1-RC with their assignments indicated.

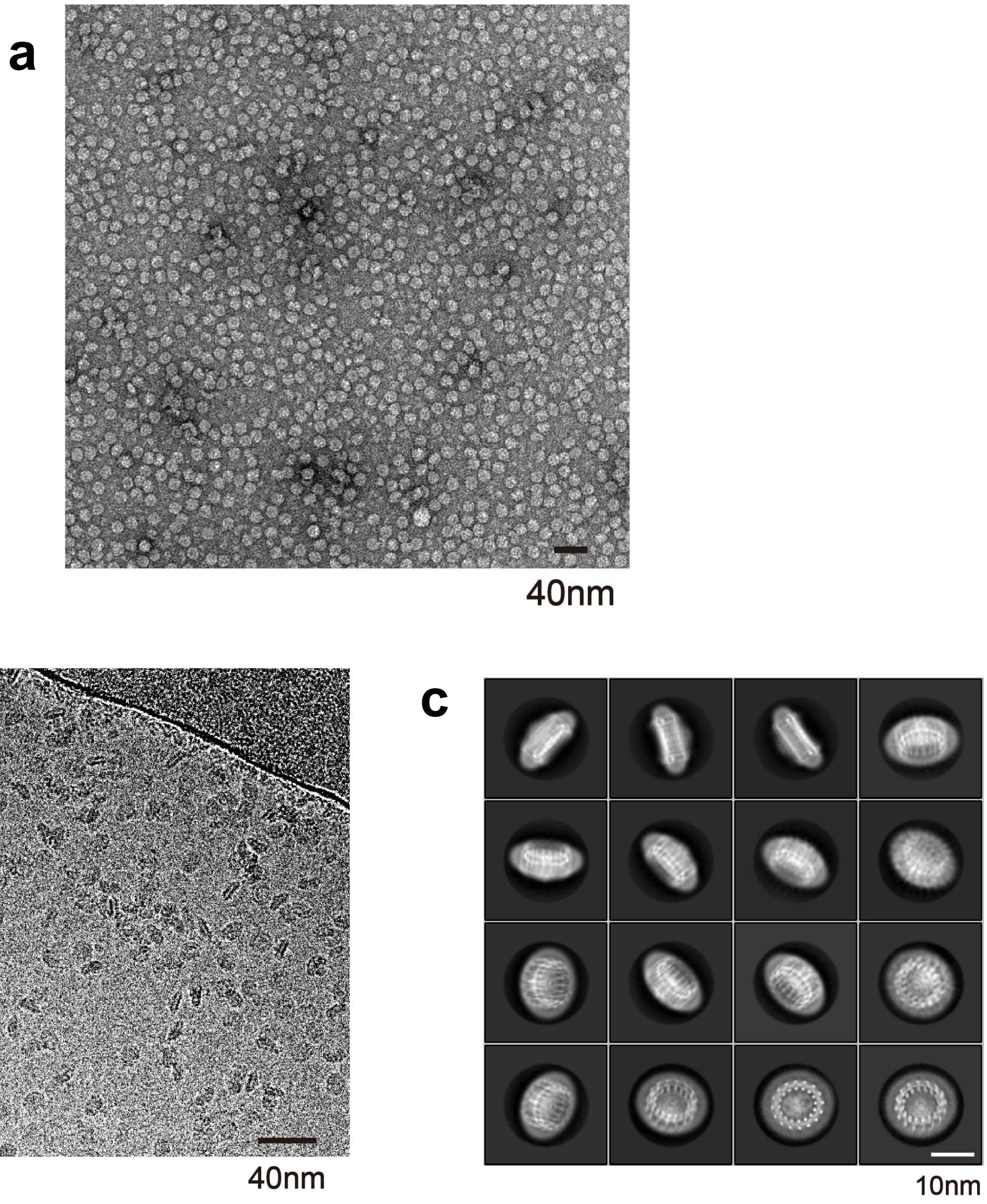


**Fig. S2 Electron micrographs of the *Rsp. rubrum* LH1-RC complex.** A representative negatively stained (**a**) and cryo-EM (**b**) micrographs of the purified LH1-RC particles. (**c**) Representative 2D class averages from cryo-EM micrographs.

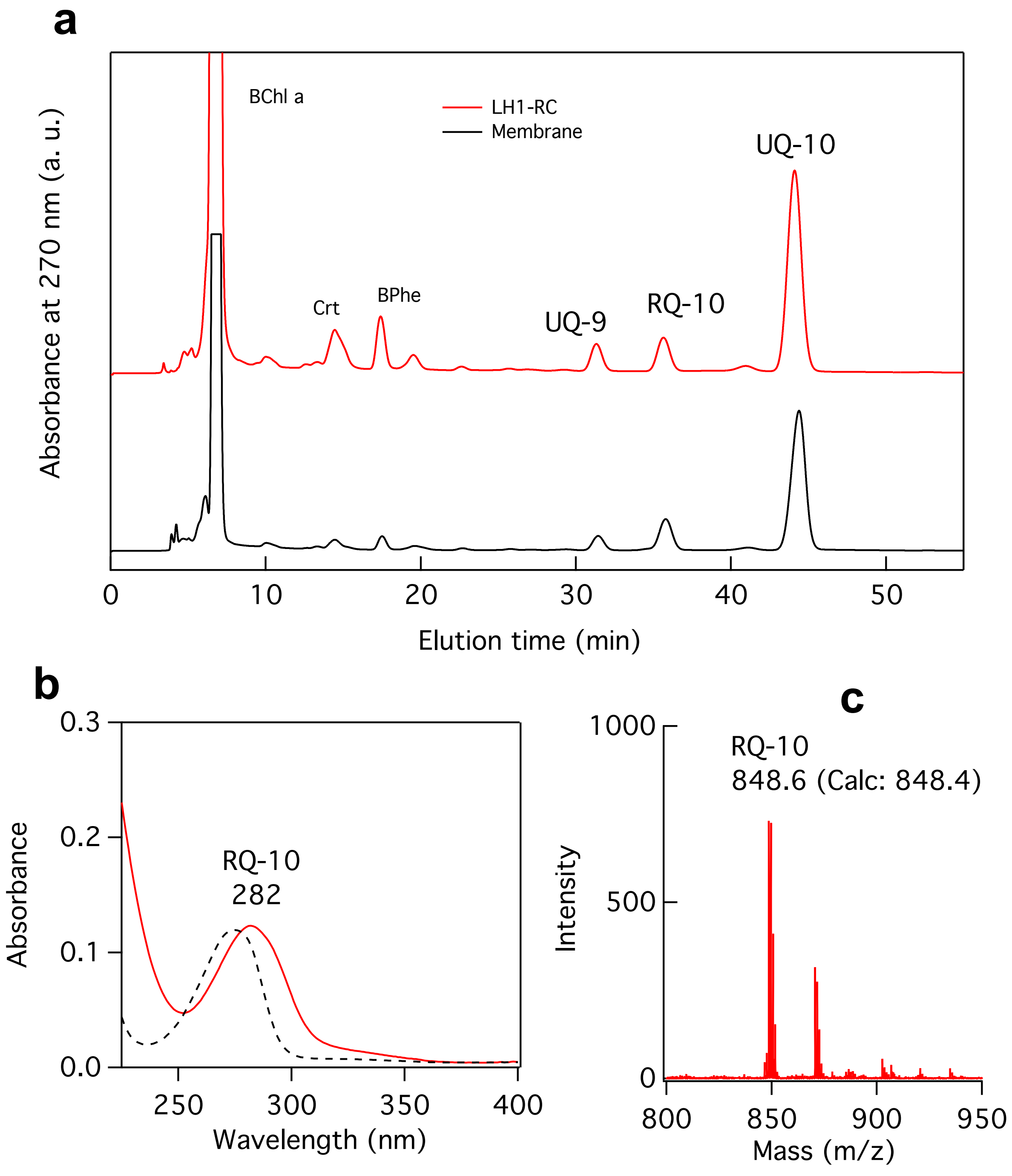


**Fig. S3 Identification of the RQ molecules in the purified LH1-RC and membranes. (a)** Reverse-phase HPLC chromatograms (TSKgel, SuperODS, 4.6×100 mm, TOSOH) of the extracted quinones and pigments isocratically eluted at 25 ˚C by 7:3 methanol/isopropanol at flow rate of 0.7 mL/min. **(b)** Absorption spectra of the RQ-10 (red curve) and UQ-10 (black dashed curve) collected from corresponding HPLC fractions in (a). **(c)** MALDI-TOF/MS spectrum of the RQ-10 collected from the HPLC fractions in (a) under the same conditions as described in Ref. 2.

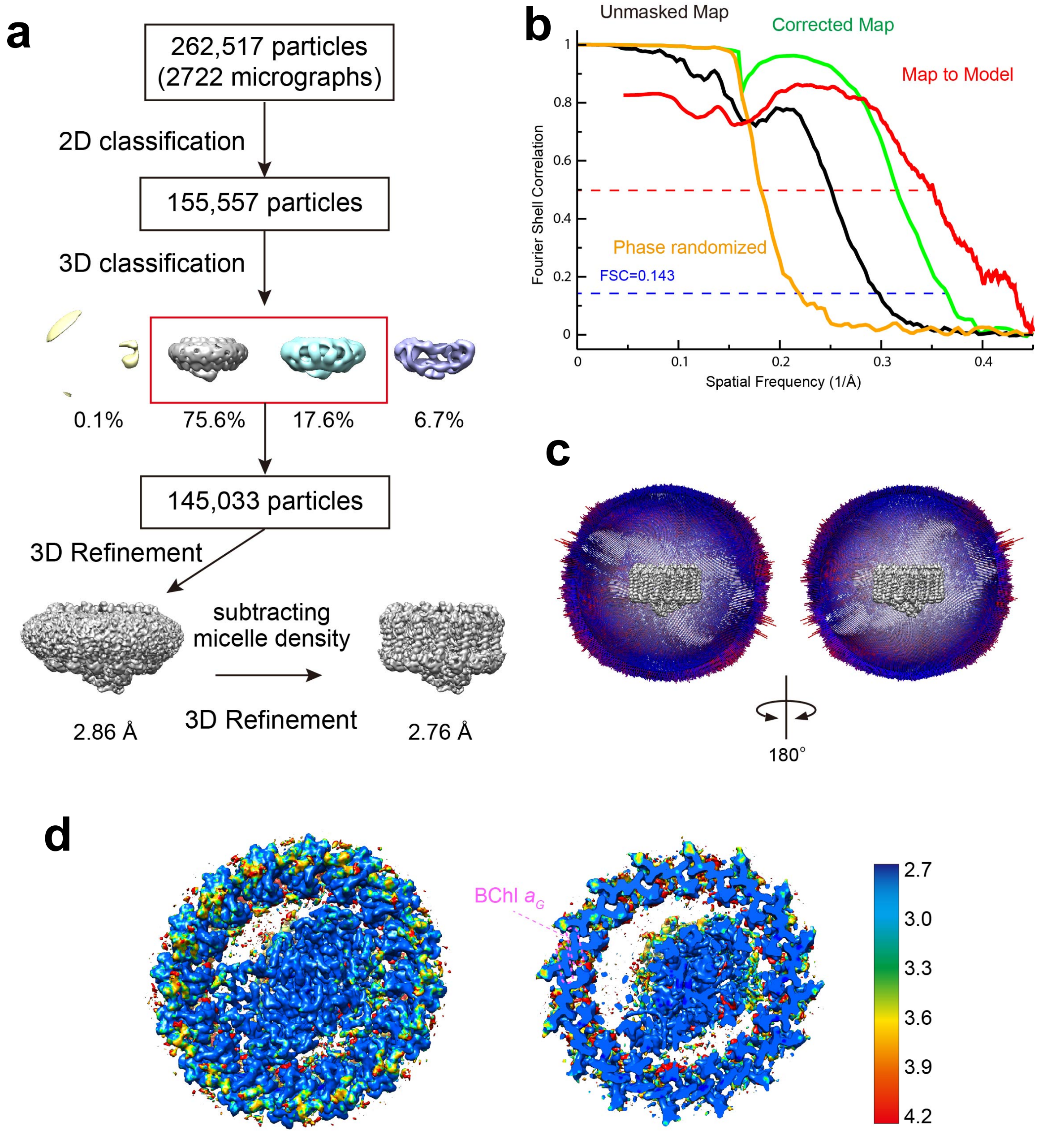


**Fig. S4 Structure determination of the *Rsp. rubrum* LH1-RC complex by cryo-EM. (a)** Image processing flow of 3D classification and reconstruction. **(b)** The Fourier shell correlation (FSC) plots of the cryo-EM map (unmasked: black, phase randomized corrected: green, phase randomized: orange) and the FSC plot of the model versus the final map (red) are superimposed. **(c)** Angular distribution of reconstructed particles. **(d)** Local resolution representation of the LH1-RC structure. Top view from periplasmic side (*left*) and a central cross sectional view (*right*). The map is shown in rainbow colors as shown in the right color bar.

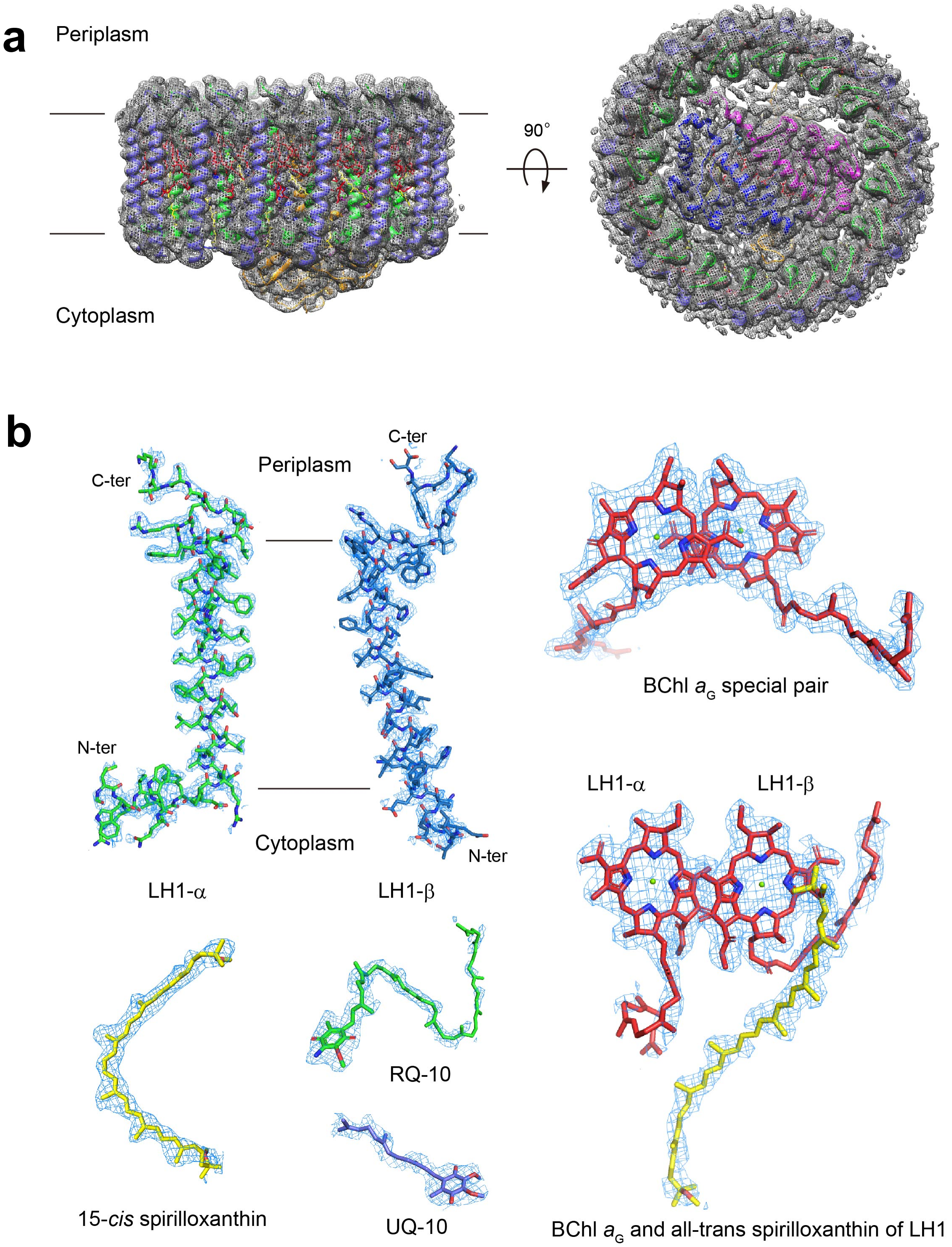


**Fig. S5 Cryo-EM densities and structural models in the *Rsp. rubrum* LH1-RC complex.** **(a)** Overall structure of the LH1-RC complex. Cartoon representation of the complex with the cryo-EM density in gray mesh. Side view (*left*) parallel to the membrane plane and top view (*right*) from periplasmic side. **(b)** Selected polypeptides and cofactors. The density maps are shown at a contour level of 4.0σ, except for the UQ-10 (2.0σ). The color codes are the same as in Fig. 1.

```
α-polypeptide
                                    10        20        30        40        50
Rsp. rubrum                 MWRIWQLFDPRQALVGLATFLFVLALLIHFILLSTERFNWLEGASTKPVQTSMVMPSSDLAV
Tch. tepidum         MFTMNANLYKIWLILDPRRVLVSIVAFQIVLGLLIHMIVLSTD-LNWLDDNIPVSYQALGKK
Alc. vinosum           MSPD-LWKIWLLVDPRRILIAVFAFLTVLGLAIHMILLSTAEFNWLEDGVPAATVQQVTPVVPQR
Trv. strain 970     MNAKSFDGMHKLWMIMNPVSTLWAIFIFQIFLGLLIHMVVLSSD-LNWHDDQIPVGYQLQGETLPVNLEMKAAQ
Rba. sphaeroides         MSKFYKIWMIFDPRRVFVAQGVFLFLLAVMIHLILLSTPSYNWLEISAAKYNRVAVAE
Rba. capsulatus          MSKFYKIWLVFDPRRVFVAQGVFLFLLAVLIHLILLSTPAFNWLTVATAHGY
Blc. viridis        MATEYRTASWKLWLILDPRRVLTALFVYLTVIALLIHFGLLSTDRLNWWEFQRGLPK
Rvi. gelatinosus            MWRIWRLFDPMRAMVAQAVFLLGLAVLIHLMLLGTNKYNWLDGAKKAPVATAVAPVPAEVTSLAQAK
Rps. palustris              MWRIWLLFDPRRALVLLFVFLFGLAIIIHFILLSTSRFNWLDGPRA
Rps. acidophila             MYKLWLLFDPRRTLVALSAFLFVLGLIIHFISLSTDRFNWLEGKPAVRA
Rps. marina                 MWKVWLLFDPRRTLVALFTFLFVLALLIHFILLSTDRFNWMQGAPTAPA

β-polypeptide
                             10        20        30        40        50
Rsp. rubrum         MAEVKQESLSGITEGEAKEFHKIFTSSILVFFGVAAFAHLLVWIWRPWVPGPNGYSALETLTQTLTYLS
Tch. tepidum           AEQKSLTGLTDDEAKEFHAIFMQSMYAWFGLVVIAHLLAWLYRPWL
Alc. vinosum           MANSSMTGLTEQEAQEFHGIFVQSMTAFFGIVVIAHILAWLWRPWL
Trv. strain 970         AEKPSTGLTESEAKEFHGLFMASMTLWFGLVVLAHILSWMYRPWL
Rba. sphaeroides     ADKSDLGYTGLTDEQAQELHSVYMSGLWPFSAVAIVAHLAVYIWRPWF
Rba. capsulatus     MADKNDLSFTGLTDEQAQELHAVYMSGLSAFIAVAVLAHLAVMIWRPWF
Blc. viridis          ADLKPSLTGLTEEEAKEFHGIFVTSTVLYLATAVIVHYLVWTARPWIAPIPKGWV
Rvi. gelatinosus     MAERKGSISGLTDDEAQEFHKFWVQGFVGFTAVAVVAHFLVWVWRPWL
Rps. palustris         MSDGSISGLSEAEAKEFHSIFVTSFFLFIVVAVVAHILAWMWRPWLPKATGY
Rps. acidophila       AEDRSSLSGVSDAEAKAFHALFVSSFTAFIVIAVLAHVLAWAWRPWIPGPKGWA
Rps. marina         AEIDRPVSLSGLTEGEAREFHGVFMTSFMVFIAVAIVAHILAWMWRPWIPGPEGYA
```

**Fig. S6 Sequence comparisons between various LH1 polypeptides with the numbering for the *Rsp. rubrum* LH1 polypeptides.** Full sequences derived from genome are shown for the *Rsp*. *rubrum* LH1 polypeptides in which the post-translationally removed residues are shown in gray fonts. The His and Trp residues that coordinate and/or hydrogen-bond to BChl *a* are shown in red, bold fonts. Residues involved in protein–protein interactions in the N- and C-terminal regions within a face-to-face subunit are shown in blue, bold fonts. Red boxes show the positions that are all aromatic residues in the α-polypeptides and rich in cationic residues in the β-polypeptides, these residue pairs may have cation–π interactions. The underlined residues in the *Rsp*. *rubrum* LH1 polypeptides form hydrophobic channels in the LH1 structure. Abbreviations: *Alc*., *Allochromatium*; *Rvi*., *Rubrivivax*.

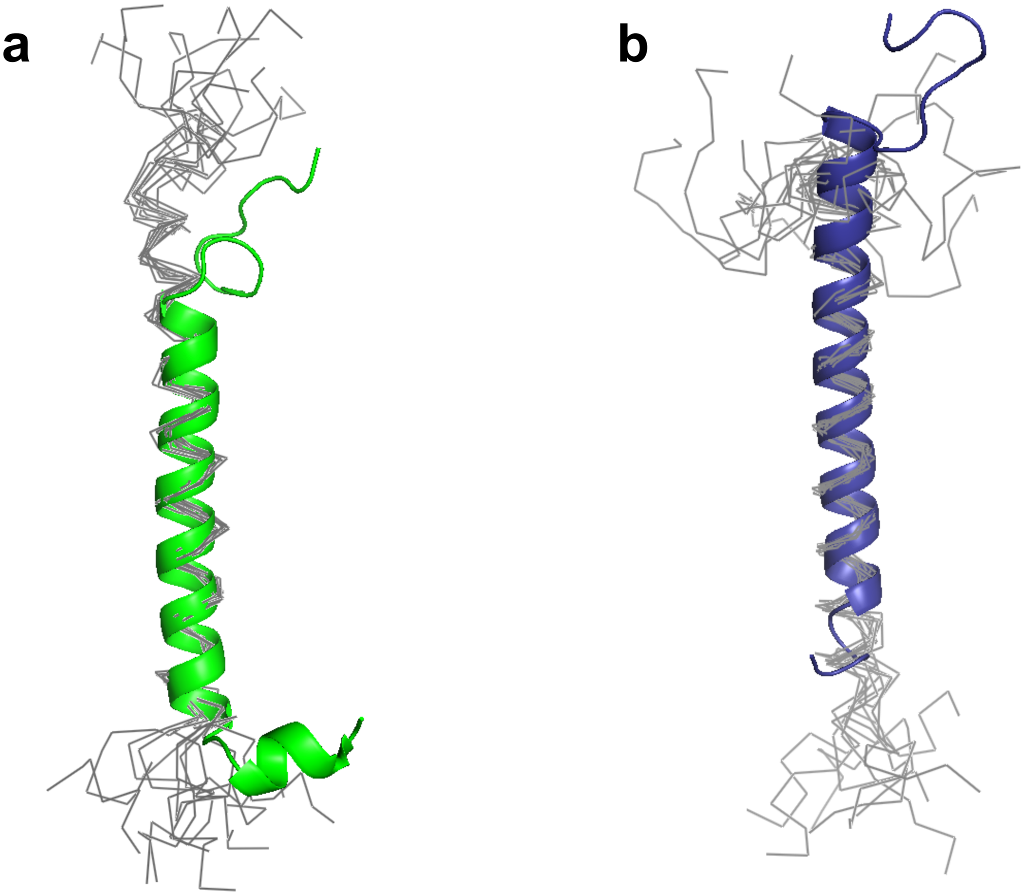


**Fig. S7 Superposition of the transmembrane domains for the structures of *Rsp. rubrum* LH1-αβ polypeptides determined by cryo-EM (this work, colored) and solution NMR (gray ensemble). (a)** LH1-α polypeptides (PDB: 1XRD for the NMR structure ensemble). **(b)** LH1-β polypeptides (PDB: 1WRG for the NMR structure ensemble).

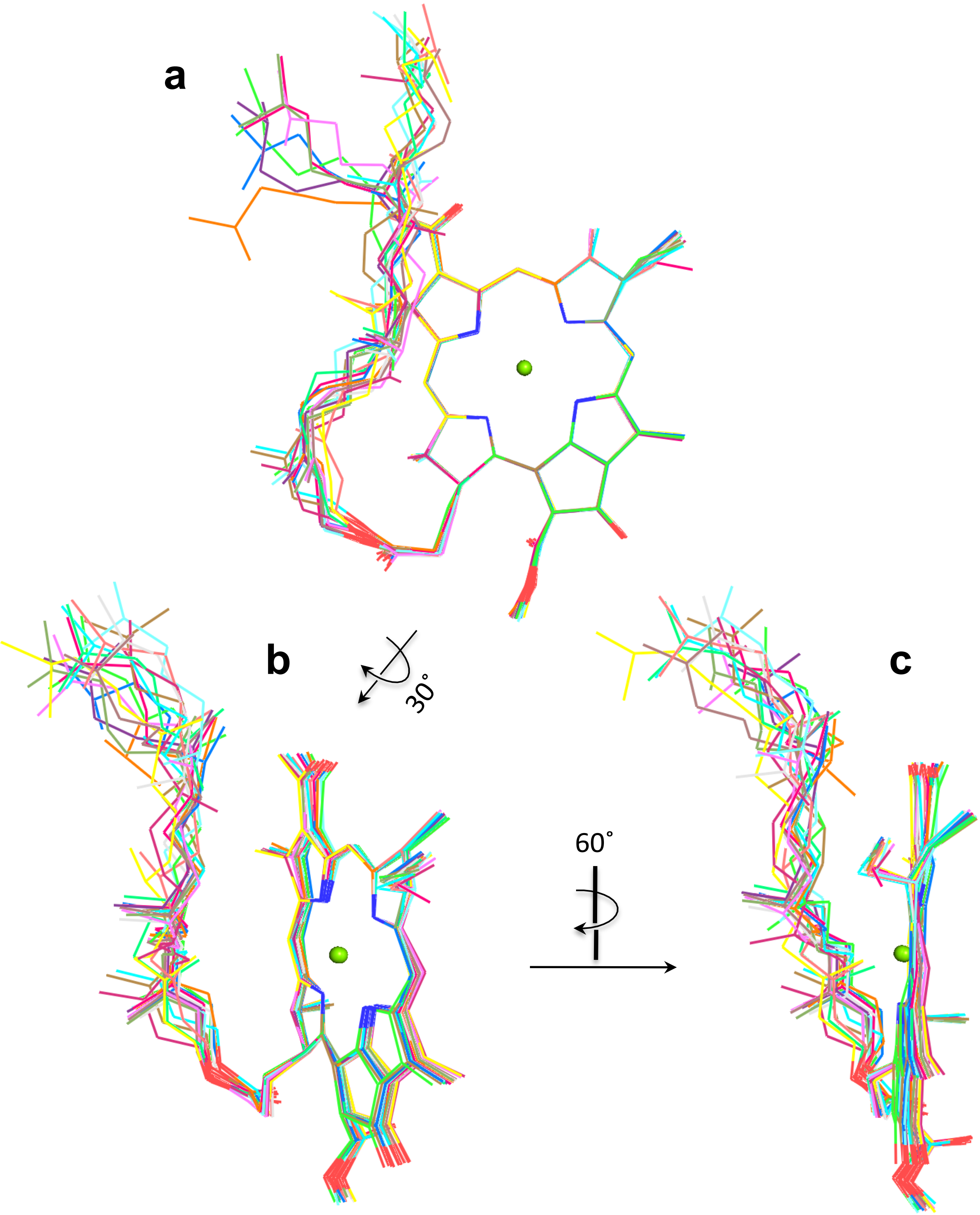


**Fig. S8 Superposition of the bacteriochlorin rings of 16 BChl *a* molecules that are esterified by phytyl group and bound to the LH1 β-polypeptides in *Tch. tepidum* (PDB: 5Y5S). (a)** Front view. **(b)** 30˚-rotated view. **(c)** 90˚-rotated view.

### RC-L subunit

```
Rsp. rubrum       MALLSFERKYRVRGGTLIGGDLFDFWVGPFYVGFFGVTTLLFTVLGTALIVWGAALGPSW  60
Rba. sphaeroides  MALLSFERKYRVPGGTLVGGNLFDFWVGPFYVGFFGVATFFFAALGIILIAWSAVLQGTW
                  ************ ****:**:****************:*::*:.**  **.*.*.*  :*

                  TFWQISINPPDVSYGLAMAPMAKGGLWQIITFSAIGAFVSWALREVEICRKLGIGYHIPF  120
                  NPQLISVYPPALEYGLGGAPLAKGGLWQIITICATGAFVSWALREVEICRKLGIGYHIPF
                  .   **: ** :.***. **:**********:.* *************************

                  AFGFAILAYVSLVVIRPVMMGAWGYGFPYGFMTHLDWVSNTGYQYANFHYNPAHMLGITL  180
                  AFAFAILAYLTLVLFRPVMMGAWGYAFPYGIWTHLDWVSNTGYTYGNFHYNPAHMIAITF
                  **.******::**::**********.****: *********** *.*********:.**:

                  FFTTCLALALHGSLILSAANPGKGEVVKGPEHENTYFQDTIGYSVGTLGIHRVGLILALS  240
                  FFTNALALALHGALVLSAANPEKGKEMRTPDHEDTFFRDLVGYSIGTLGIHRLGLLLSLS
                  ***..*******:*:****** **: :: *:**:*:*:* :***:*******:**:*:**

                  AVVWSIICMILSGPIYTGSWPDWWLWWQKLPFWNHG------
                  AVFFSALCMIITGTIWFDQWVDWWQWWVKLPWWANIPGGING
                  **.:* :***::*.*: ..* *** ** ***:* :
```

### RC-M subunit

```
Rsp. rubrum       MSEYQNILTGVQVR-TAPHSAPIAKGIFPRLGKPGFSYWLGKIGDAQIGPIYLGTTGVLS  59
Rba. sphaeroides  MAEYQNIFTQVQVRGPADLGMTEDVNLANRSGVGPFSTLLGWFGNAQLGPIYLGSLGVLS
                  *:*****:* **** .*  . .   .:  * *   **  ** :*:**:******: ****

                  LVFGFFAIEIIGFNLLASVNWSPMEFGRQFFWLGLEPPAAEYGLGFAP-LAEGGWWQIAG  118
                  LFSGLMWFFTIGIWFWYQAGWNPAVFLRDLFFFSLEPPAPEYGLSFAAPLKEGGLWLIAS
                  *. *:: :  **: :  ...*.*  * *::*::.*****.****.**. * *** * **.

                  FFLTTSILLWWVRMYRRARALKMGTHTAWAFASAIFLFLSLGFIRPLLMGNFSESVPFGI  178
                  FFMFVAVWSWWGRTYLRAQALGMGKHTAWAFLSAIWLWMVLGFIRPILMGSWSEAVPYGI
                  **: .::  ** * * **:** **.****** ***:*:: ******:***.:**:**:**

                  FPHLEWTNSFSLNYGNFFYNPFHMLSIAFLYGSALLFAMHGATILAVSRLGGDREVEQIT  238
                  FSHLDWTNNFSLVHGNLFYNPFHGLSIAFLYGSALLFAMHGATILAVSRFGGERELEQIA
                  *.**:***.*** :**:****** *************************:**:**:***:

                  DRGTAAERAALFWRWTMGFNATMESIHRWAWWFAVLCTFTGAIGILLTGTVVDNWFEWGV  298
                  DRGTAAERAALFWRWTMGFNATMEGIHRWAIWMAVLVTLTGGIGILLSGTVVDNWYVWGQ
                  ************************.***** *:*** *:**.*****:*******: **

                  KHGLAPAP
                  NHGMAPLN
                  :**:**
```

**Fig. S9 Sequence comparisons of L- and M-subunits in the RCs between *Rsp. rubrum* and *Rba sphaeroides*.** The His residues that coordinate and/or hydrogen-bond to special pair BChls *a* are shown in red, bold fonts. Residues involved in interactions with the RQ-10 head group and Fe atom are shown in blue, bold fonts.

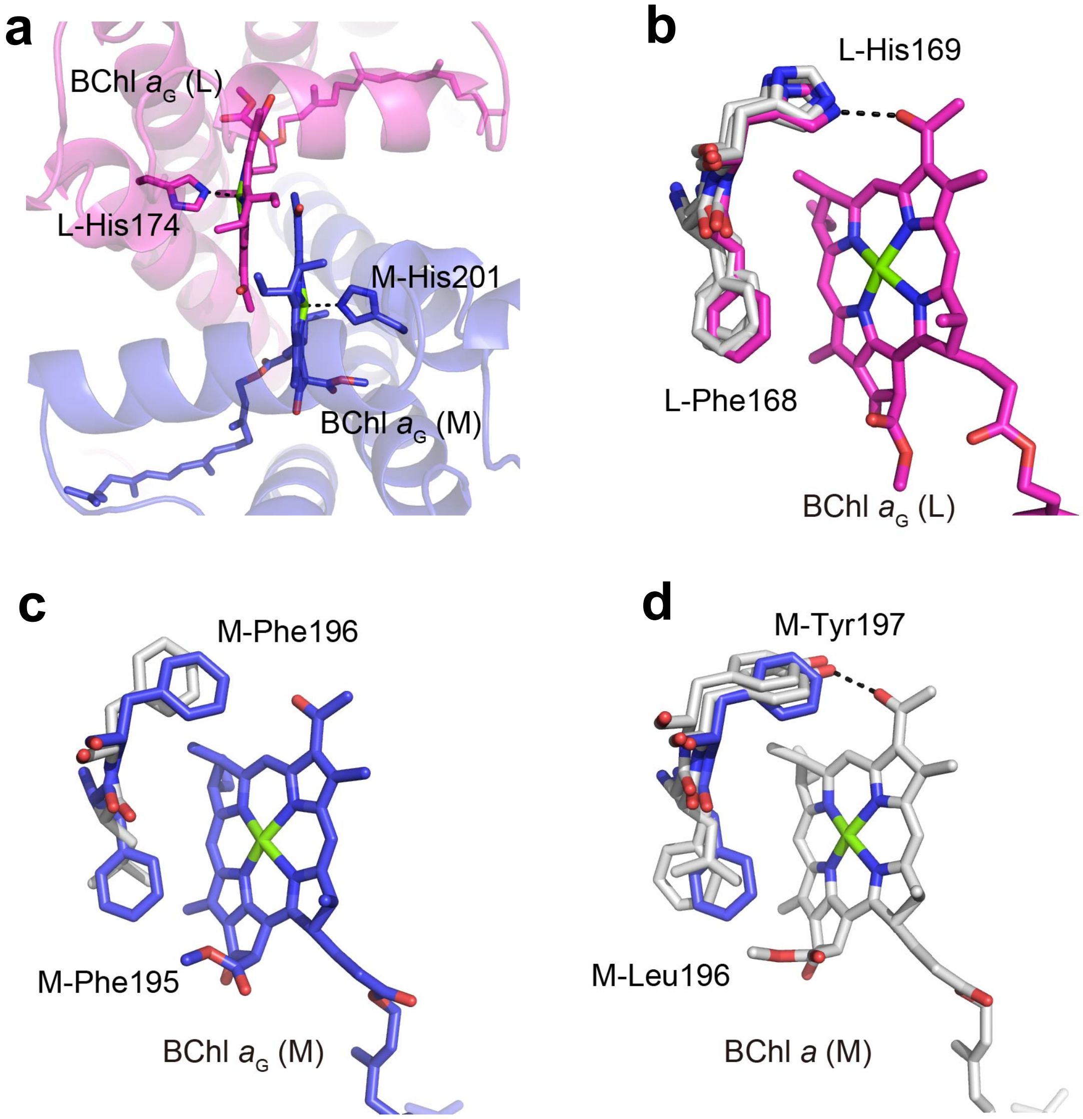


**Fig. S10 Key residues around the RC special pair BChl *a* dimer of *Rsp. rubrum*.** **(a)** Top view of the BChl $a_G$ dimer from periplasmic side. Central Mg atoms of the BChl $a_G$ are coordinated by the His residues in the L- and M-subunits. Color scheme: L-subunit, magenta; M-subunit, blue. **(b)** Superposition of the residues in the RC L-subunits of *Rsp. rubrum* (magenta), *Tch. tepidum* (gray), *Trv.* strain 970 (gray), and *Rba. sphaeroides* (gray) that form hydrogen bond or interact with the BChl *a* molecule. **(c)** Superposition of the residues in the RC M-subunits of *Rsp. rubrum* (blue) and *Rba. sphaeroides* (gray) that interact with the BChl *a* molecule. **(d)** Superposition of the residues in the RC M-subunits of *Rsp. rubrum* (blue), *Tch. tepidum* (gray), and *Trv.* strain 970 (gray) that form hydrogen bond or interact with the BChl *a* molecule. A hydrogen bond is shown as a black dotted line. Only this panel, amino acid labeling of M-subunit of *Tch. tepidum*.

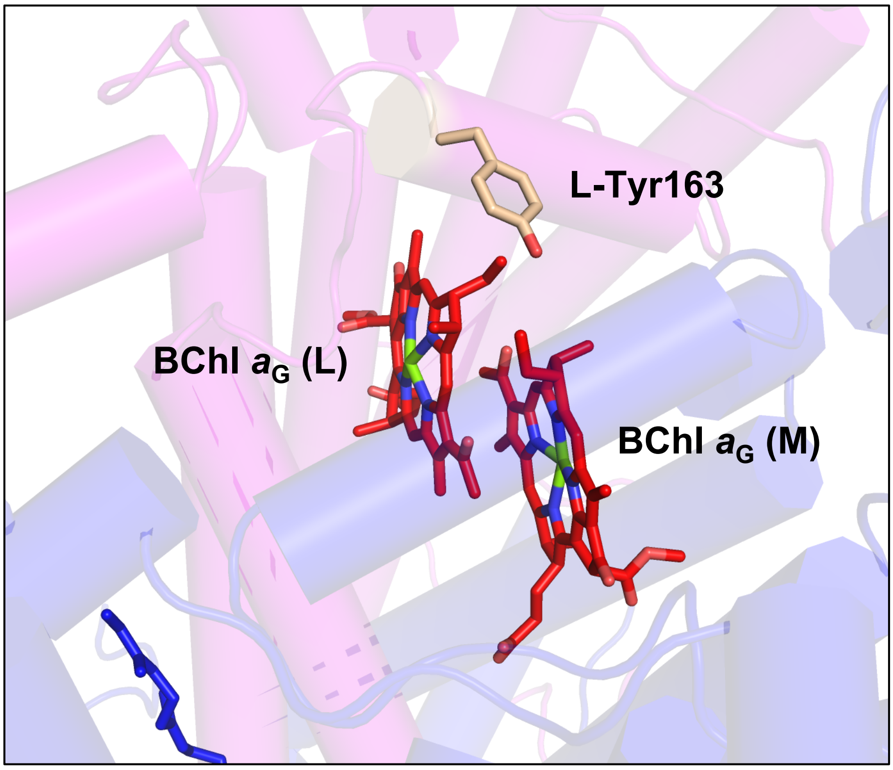


**Fig. S11 A conserved Tyr residue positioned on the periplasmic surface in close proximity to the special pair BChls $a_G$.** This Tyr is considered to serve as a bridging residue for the electron transfer from soluble cytochrome $c_2$ to the RC special pair.

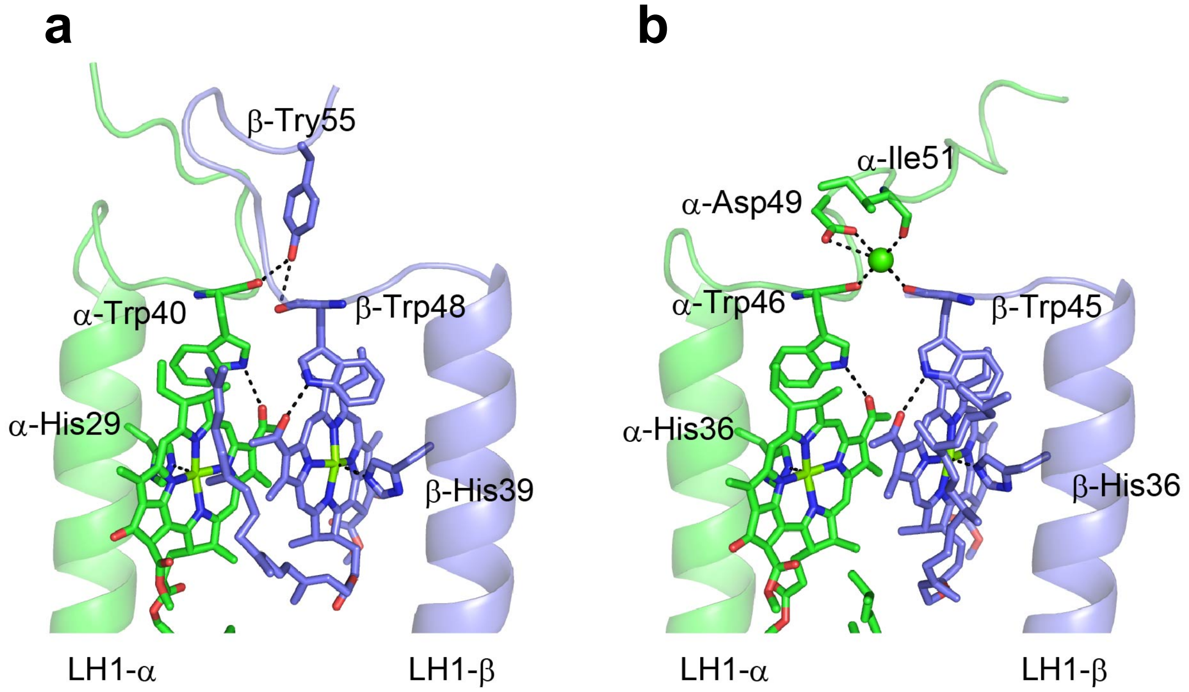


**Fig. S12 Comparison of back-to-back subunit structures from *Rsp. rubrum* and *Tch. tepidum*.** (**a**) LH1-αβ polypeptides of *Rsp. rubrum*. (**b**) LH1-αβ polypeptides of *Tch. tepidum*. Hydrogen bonds or salt bridges are indicated as black dotted lines. Color scheme: LH1-α, green; LH1-β, slate blue; $Ca^{2+}$, green ball.
